# Cascading consequences of shifting ice phenology in an exploited lake

**DOI:** 10.1101/2024.10.11.617776

**Authors:** Daisuke Goto, Erin S. Dunlop, Joelle D. Young, Donald A. Jackson

## Abstract

Ice phenology (onset and breakup of ice cover) regulates the timings of seasonal life history events of many animal and plant species in high-latitude lakes in the Northern Hemisphere, promoting the coexistence of species that share resources like food and shelter. Increasingly warmer, more variable Earth’s climate is, however, shifting ice phenology, modifying habitat conditions, and in turn reshaping the population and interaction dynamics of many species, threatening their persistence. Applying multivariate autoregressive state-space time series modeling to extensive historical and contemporary monitoring records (spanning 36 to 113 years) of limnological and biological surveys and fishery catches, we explore how shifting ice phenology propagates through a food web over time by modulating thermal profiles and limnological properties in Lake Simcoe, a large, intensely exploited system in southern Ontario, Canada. Analysis shows that increasingly earlier ice breakups, later ice freeze-ups, earlier thermal stratification onset, and later turnover since the early 1980s, attenuated algal production. This in turn, combined with increasingly more abundant invasive species (dreissenids) that promoted greater water clarity over prolonged stratification periods, diminished zooplankton abundances. These limnological changes filtered through plankton communities further modified the population and community dynamics of higher trophic-level species like fish with differential thermal preferences, interactively with time-varying non-climate drivers. Although management interventions–reductions in nutrient loading and fishing effort in particular–in recent decades aided the recoveries of depleted cold-adapted fish populations (including lake trout, lake whitefish, and cisco), their interaction patterns and strengths with species adapted to warmer conditions were reshuffled by advanced timings of ice breakup and delayed timings of ice freeze-up. These findings reveal how complex food web dynamics can emerge from ecologically divergent responses that reshape species interactions within and among trophic levels under shifting ice phenology filtered through bottom-up processes, likely buffering destabilizing effects of a changing climate.

## INTRODUCTION

Life history events of wild animal and plant populations (timings of migration, flowering, etc.) are often synchronized with periodicity or seasonality in physical environments, driving their population and community dynamics in marine (Poloczanska et al. 2013), terrestrial (Piao et al. 2019), and freshwater (Woods et al. 2022) systems. Increasingly warmer Earth’s climate is, however, disrupting and reshaping these patterns ubiquitous across biomes (Visser and Both 2005, Post 2013, Thackeray et al. 2016, Cohen et al. 2018). Warming rates in lakes are especially greater compared to those in marine and terrestrial ecosystems (O’Reilly et al. 2015). Existing evidence indicates that these trends of accelerated warming (and more frequent thermal extremes, Woolway et al. 2021a) in lakes will continue in many parts of the world (Woolway and Maberly 2020), modifying their physical and chemical properties and triggering shifts in habitat condition that may diminish primary and secondary productivity (O’Reilly et al. 2003, Kraemer et al. 2021).

Warming surface water, combined with slowed wind speed, can modulate the seasonality of limnological events like timings of ice breakup in spring and freeze-up in autumn especially in high-latitude systems in the Northern Hemisphere (Benson et al. 2012, Sharma et al. 2019, Woolway et al. 2021b), triggering cross-seasonal domino effects (Hampton et al. 2017, Hébert et al. 2021). Past research based on ice phenology records spanning several centuries shows that ice breakup dates advanced on average by 11 days per century, whereas ice onset dates delayed by 7 days, shortening ice cover duration by 17 (Magnuson et al. 2000, Sharma et al. 2021b). Some of these lakes could lose winter ice cover completely by the end of the twenty first century (Sharma et al. 2019, Sharma et al. 2021a). Winter ice cover regulates timings of life history events (before or after the period), demography, and population dynamics of many species in northern temperate lakes (Shuter et al. 2012, Hampton et al. 2017, McMeans et al. 2020). Changes in the cues that trigger seasonal life history events can be further advanced or delayed by local physical environmental conditions (Thackeray et al. 2016). Winter ice cover periods thus may act as a mediator to promote coexistence of species that compete for shared resources like food and shelter (McMeans et al. 2020).

Winter ice cover duration may determine temporal habitat partitioning and interaction strength among mobile, higher trophic-level species with varying thermal preferences like fish, likely reshaping their community structure (Helland et al. 2011, McMeans et al. 2020). Changes in differential seasonal habitat use by these vertebrate consumers triggered by shortened ice cover duration and milder winter conditions as climate warms may force competing species to share habitat and its limited resources, compromising survival and reproduction of competitively inferior species (Shuter et al. 2012, Hovel et al. 2017, McMeans et al. 2020) and perhaps population persistence. Increasingly earlier ice breakup in spring can, for example, alter thermal habitat and shorten the duration of energy-rich littoral habitat use by cold-adapted (cold-water) species like salmonids in warming lakes, thereby reducing their fitness (Guzzo et al. 2017, Caldwell et al. 2020) while expanding habitat size for warm-water species like smallmouth bass (Sharma et al. 2007). Cold-water species may be squeezed out their habitat directly (via thermal stress, Guzzo et al. 2017, Caldwell et al. 2020) and indirectly (via amplified competition with warm-water species, Helland et al. 2011, Shuter et al. 2012, McMeans et al. 2020) as ice phenology shifts under climate warming.

Shifting ice phenology in northern temperate lakes also can disrupt well-defined seasonal cycles of pelagic algae, which form the base of pelagic food webs (Adrian et al. 1999, Michelutti et al. 2020). Shorter winter ice cover resulting from delayed onset and advanced breakup (longer growing season) can promote primary production during ice-free periods (Hébert et al. 2021). Longer and more intense thermal stratification in the growing season can further modify vertical habitat use of pelagic species and their interaction strengths (Francis et al. 2011). Varying responses to shifting timings of primary production can emerge as complex, food web restructuring confounded with other time-varying limnological processes, propagating through species interactions (Winder and Schindler 2004b, Schindler et al. 2005, Hébert et al. 2021). Although climate warming may to some extent promote primary production and in turn increase fat content in some primary consumers like copepods (Hébert et al. 2021), asynchronized shifts in life history phenology among trophic levels may result in trophic mismatches (Winder and Schindler 2004a, Jan et al. 2024) and thus disruption in trophic networks (Woodward et al. 2010, Hébert et al. 2021), likely triggering cascading effects on system-wide food web dynamics.

Here we report an empirical analysis to explore how shifting ice phenology propagates through a food web from primary producers to top predators over time by modulating thermal profiles and limnological properties in Lake Simcoe, southern Ontario, Canada (Fig. 1a). Lake Simcoe is a large, intensely exploited system exposed to a variety of human-induced pressures including excessive fishing (Dunlop et al. 2019), excessive nutrient loading (Evans et al. 1996), and exotic species introduction (North et al. 2013) over more than a century, resulting in trophic cascades and regime shifts (Goto et al. 2020). In recent decades the lake also began experiencing climate warming-induced shifts in ice phenology and thermal stratification timings, and changes in other limnological properties (Fig. 1c–e). These climate change-induced shifts would also influence the population dynamics and interactions of species that depend on these timings as cues for their seasonal life history events (Shuter et al. 2012). Applying time series modeling to extensive monitoring records of ice phenology, limnological and biological surveys, and fishery catches from the lake, we explore a) how shifting lake phenology reshapes fish community dynamics interactively with existing pressures from human activities, and b) how bottom-up processes modified by lake phenology propagate through food web networks under a changing climate.

**Figure 1.**
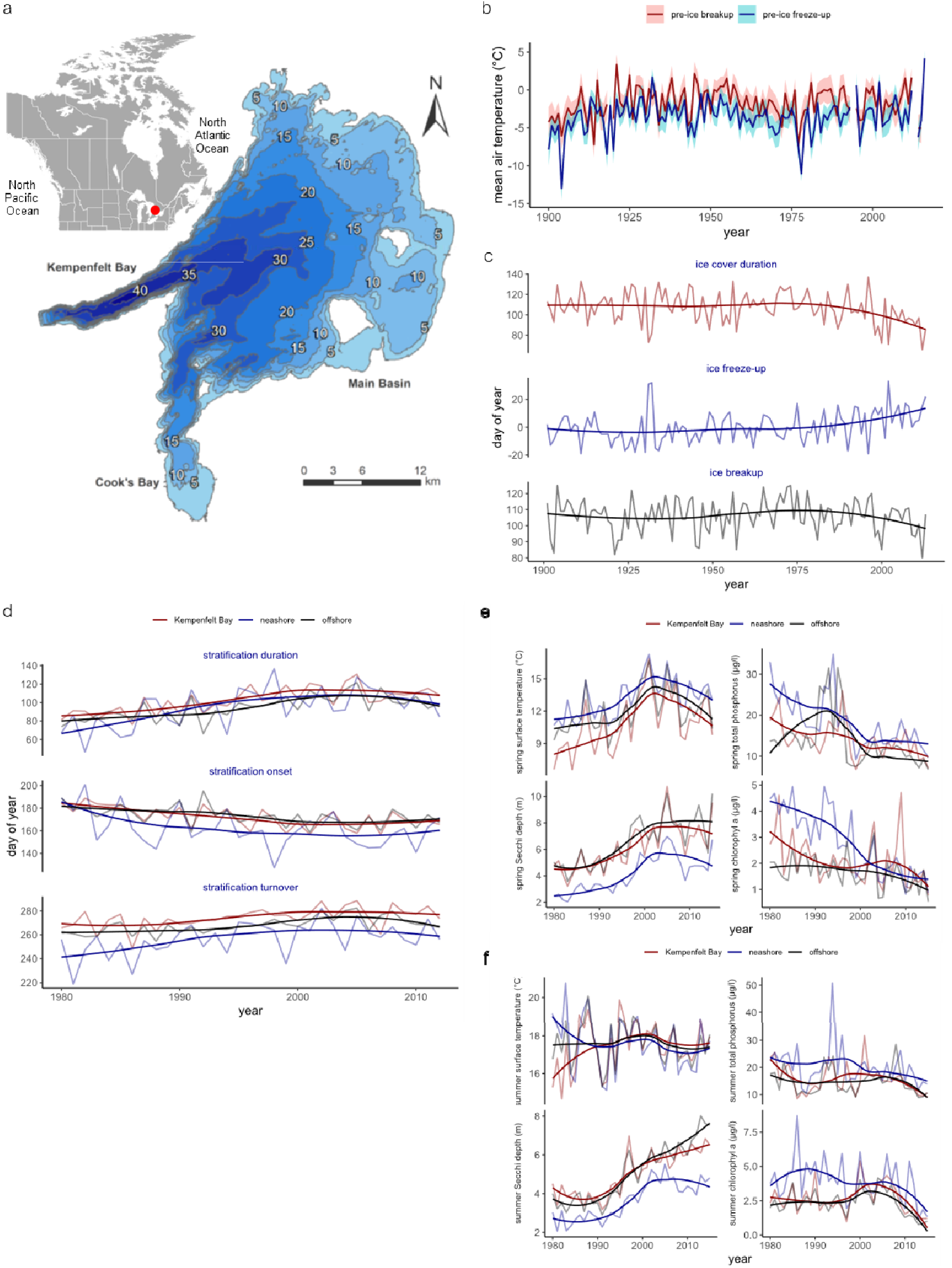
Climate, ice phenology, and limnological conditions of a study system, Lake Simcoe. (a) Lake bathymetry, (b) daily mean air temperatures two months before ice breakup and freeze-up during 1900–2015, (c) ice phenology (ice cover duration, ice freeze-up, and ice breakup) during 1901–2013, (d) thermal stratification phenology (duration, onset, and turnover) in Kempenfelt Bay and offshore and nearshore areas during 1980–2015, and (e) spring and (f) surface water temperature, Secchi depth, total phosphorus concentration, and chlorophyl *a* concentration in Kempenfelt Bay and offshore and nearshore areas during 1980–2015.

## METHODS

To investigate time-varying food web responses to shifting ice phenology in Lake Simcoe, we applied a series of multivariate autoregressive state-space (MARSS) models to time series of fish and plankton communities along with climate and limnological covariates. In the following we describe (a) reconstruction of fish community dynamics by integrating long-term historical and contemporary monitoring and catch data, followed by analyses to test (b) regional and local climate effects on shifting ice phenology and its effects on thermal stratification phenology, (c) effects of shifting ice phenology and limnological changes on algal production, (d) interactive effects of shifting ice phenology and long-term human activities on fish community dynamics, and (e) propagating bottom-up effects of shifting ice phenology on food web dynamics.

### Data

We compiled time series data of fish abundance indices (catch per unit effort (per hour) or CPUE) collected primarily through five historical and modern annual scientific monitoring and creel surveys programs conducted in Lake Simcoe since 1951 by the Lake Simcoe Fisheries Assessment Unit of Ontario Ministry of Natural Resources and Forestry (MNRF); fall index trap netting (FITN) survey (1951–2006), winter creel survey (1961–2015), summer creel survey (1965–2015), nearshore community index netting (NSCIN) survey (1992–2015), and offshore benthic gillnet index netting (OSBIN) survey (2003–2015). The FITN survey was conducted with 3.05-m trapnets targeting primarily cold-water species spawning at fixed locations (North Georgian Island and Strawberry Island) from late September to early December (MacRae 2001). The winter creel survey was conducted lake-wide through angler interviews (collecting effort (hours spent by anglers), catch, harvest, and auxiliary information like body size) over 50 continuous days from late January to mid-March (ice-covered period) using a stratified roving method (MacRae 2001). The same method was also applied to the summer creel survey conducted lake-wide from early May to September (MacRae 2001). The NSCIN survey is a standardized sampling program using 1.8-m trapnets set at 30 randomly selected sites (22 hours per sampling occasion) targeting primarily cool-and warmwater species between late summer and early autumn (Robillard 2010). Likewise, the OSBIN survey is another standardized sampling program using gillnets (137.2 m × 2.4 m with monofilament mesh) set at ∼30 randomly selected offshore locations (>25 m) targeting primarily cold-water species during mid-summer (Dolson 2012). For analysis we excluded species caught in less than 10 years in each survey or species caught in only one survey type. In Lake Simcoe yearlings or fingerings of lake trout and lake whitefish have been stocked since the late 1800s to support commercial and then recreational fisheries (Dunlop et al. 2019). In this study CPUE timeseries of the wild and stocked components (both in fishery-independent and -dependent data) were combined because the main focus of this study was to quantify the strength of species interactions and the roles of climate change-induced changes in limnological properties in the interactions. The final data set comprises 75 time series spanning 65 years (1951–2015) of 19 numerically dominant species with differential thermal preferences and spawning timings, representing the fish community in the lake (Table 1).

**Table 1.** Study fish species in Lake Simcoe and their thermal preferences and spawning seasons.

| common name | scientific name | thermal preference | spawning season |
| --- | --- | --- | --- |
| black crappie | *Pomoxis nigromaculatus* | warm | spring |
| bluegill | *Lepomis macrochirus* | warm | spring |
| bowfin | *Amia calva* | warm | spring |
| brown bullhead | *Ameiurus nebulosus* | warm | spring |
| burbot | *Lota lota* | cold | autumn |
| channel catfish | *Ictalurus punctatus* | cold | autumn |
| cisco | *Coregonus artedii* | cold | autumn |
| common carp | *Cyprinus carpio* | warm | spring |
| lake trout | *Salvelinus namaycush* | cold | autumn |
| lake whitefish | *Coregonus clupeaformis* | cold | autumn |
| largemouth bass | *Micropterus nigricans* | warm | spring |
| northern pike | *Esox lucius* | warm | spring |
| pumpkinseed | *Lepomis gibbosus* | warm | spring |
| rainbow smelt | *Osmerus mordax* | warm | spring |
| rock bass | *Ambloplites rupestris* | warm | spring |
| smallmouth bass | *Micropterus dolomieu* | warm | spring |
| walleye | *Sander vitreus* | cool | spring |
| white sucker | *Catostomus commersonii* | warm | spring |
| yellow perch | *Perca flavescens* | cool | spring |

We compiled time series data of limnological properties and plankton abundances collected through a variety of historical and modern annual monitoring survey programs, as well as local and regional climate indices from databases. Ice phenological records (ice breakup, freeze-up, and duration) have been collected annually since 1901. Ice phenology of many temperate lakes in the Northern Hemisphere is correlated with regional climate signals like North Atlantic Oscillation (NAO) and local air temperature (Straile et al. 2003, Mishra et al. 2011). For regional climate indices, monthly time series of the NAO index and the Atlantic Multidecadal Oscillation (AMO) index were downloaded (computed from the extended reconstructed SST (ERSSTv5, NOAA Physical Sciences Laboratory; https://psl.noaa.gov) and averaged over winter months (when the atmosphere is most active in the region, (Stenseth et al. 2003). Because the temporal coverage of air temperature from the weather station at Lake Simcoe (44°26′12″N at 79°20′21″W) was limited, we compiled historical records (1901–2015) of daily air temperature from weather stations within a 50 km radius downloaded from Environment and Climate Canada (https://climate.weather.gc.ca/). Temperature data were averaged across the stations and then across the months leading up to ice freeze-up (November to January in the following year) and breakup (February to April) (Fig. 1b).

We obtained zooplankton community and limnological data from long-term monitoring survey programs conducted by the Ontario Ministry of the Environment, Conservation and Parks (MECP) during 1980–2015. Zooplankton densities were used as an indicator as the food sources of planktivorous fish species, including the early life stages of benthivores and piscivores before switching to zoobenthos and fishes, filtering limnological changes under ice phenological shifts. Detailed sampling protocols were documented previously (Nicholls and Tudorancea 2001). Briefly, plankton samples (number l^-1^) were collected with an 80-μm mesh net biweekly between mid-April and late November (ice free season) at nine fixed stations (three at the main basin (offshore), three nearshore areas, and three in Kempenfelt Bay) in the lake, where limnological measurements were also taken. In this study we selected epilimnion (surface), total phosphorous concentration (µg l^-1^), Sacchi depth (an indicator of water clarity, m), and chlorophyl *a* concentration (as a proxy for algal abundance, µg l^-1^). For analysis we aggregated zooplankton data to the genus level. Stratification phenology (onset, turnover, and duration) was also estimated from water temperature data at two offshore and one Kempenfelt Bay stations.

### Time series analysis

To explore how warming climate-induced shifts in ice phenology, ecosystem processes, and food web dynamics in Lake Simcoe along with continued human perturbations, we applied multivariate autoregressive state-space (MARSS) modeling (Holmes et al. 2023) to the compiled time series data to account for process and observation uncertainties associated with modeled state variables. We first reconstructed fish community dynamics, followed by testing effects of climate change on ice phenology, 2) ice phenological and limnological changes on algal abundance, 3) ice phenological changes and human activities on fish community dynamics, and 4) ice phenological and limnological changes on food web dynamics. Specific structures of MARSS models applied to different datasets are described below.

### Reconstruction of fish community dynamics

Historical and contemporary fish monitoring and catch surveys varied in spatial and temporal coverage with all species included in analysis caught in multiple surveys with overlapping spatial and temporal coverages. To reconstruct fish community dynamics we first estimated relative abundances by integrating survey time series per species using first-order MARSS modeling (Tolimieri et al. 2017, Zhu et al. 2018, Rook et al. 2022). Here the model consists of two main equations that describe the process and observation components of the state, fish abundance, as follows

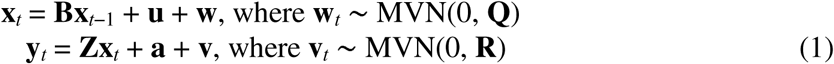

where **x***_t_* is an *n* × 1 matrix of the log-transformed true (unobserved) abundances of *n* species in year *t*, **B** is an *n* × *n* identity matrix (1s on the diagonal and 0s elsewhere), **u** is an *n* × 1 matrix of the long-term mean population growth rates (the process component is in effect the Gompertz model, (Gompertz 1825) of *n* species, **w***_t_* is the process error that follows a multivariate normal distribution with mean 0 and variance-covariance matrix **Q**, **y***_t_* is an *m* × 1 matrix of the log-transformed observed abundances (CPUEs) in year *t* (all zeros in fishery-independent times series were treated missing and estimated in model-fitting), **Z** is an *n* × *m* design matrix that defines how *m* time series of CPUEs are linked to *n* species, **a** is an *m* × 1 matrix of scaling parameters that defines how CPUEs from different surveys are combined per species, and **v***_t_*is the observation error that follows a multivariate normal distribution with mean 0 and variance-covariance matrix **R**. Because of differences in scale among surveys, we standardized the CPUE time series (z-scores) prior to model-fitting. In estimating the **a** matrix all species-specific time series were scaled to the contemporary scientific monitoring (OBSIN or NSCIN) surveys because they provide least biased estimates of lake-wide relative abundance. We selected the variance-covariance structures of **Q** and **R** based on the model fitted to 12 select structures (Table S1) fitted with a maximum likelihood (ML) method with an Expectation–Maximization algorithm (Holmes et al. 2023). These structures were constructed based on assumptions of data source and population biology using combinations of equal process error for all species, species-specific process error without and with temporal correlation, equal observation error for all time series, equal observation error per species, equal variances between the observed abundances are independent (no covariance in **R**). The model selection was based on the lowest Akaike information criterion (AICc) score corrected for small sample size. AICc was computed as AICc = –2ln(*L*) + 2*k*(*k* + 1)/(*n* – *k* – 1), where *L* is the likelihood of the model, *k* is the number of parameters, and *n* is the sample size. Confidence intervals were estimated by parametric bootstrapping (*n* = 500) (Hampton et al. 2006).

### Climate effects on shifting ice phenology and thermal profiles

We quantified the role of changing climate in lake ice phenology and stratification shifts using MARSS modeling in two steps: testing effects of 1) regional climate and local air temperatures in late autumn and early spring (the periods leading up to ice freeze-up and breakup) near the lake on ice phenology and 2) ice phenological changes on stratification timings as

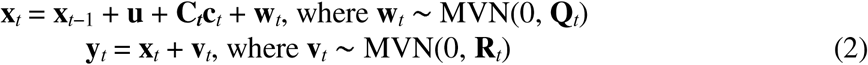

where **x***_t_* is the matrix of the true (unobserved) ice phenology or stratification timing (freeze-up/onset, breakup/turnover, and duration) in year *t*, **u** is the vector of parameters, **C*_t_*** is the matrix of climate effects on ice phenology or ice phenology effects on stratification in year *t*, **c***_t_* is the matrix of the covariates in year *t*, **w***_t_* is the process error that follows a multivariate normal distribution with mean 0 and variance-covariance matrix **Q**, **y***_t_* is the matrix of the observed ice phenology or stratification timing in year *t*, and **v***_t_* is the observation error that follows a multivariate normal distribution with mean 0 and variance-covariance matrix **R**. The stratification timings were recorded in three areas of the lake with varying depths (nearshore, offshore, and Kempenfelt Bay–a deepest part of the lake, Fig. 1) and thus modeled separated (as a 9 × 1 matrix for **x***_t_*). The model selection was based on the lowest AICc score. Confidence intervals were estimated by parametric bootstrapping (*n* = 500) (Hampton et al. 2006).

### Effects of lake phenological and limnological changes on seasonal algal blooms

We quantified effects of shifting ice phenology and thermal stratification timing on algal blooms using MARSS modeling with the same model structure as eqn. 2, except **x***_t_* being the matrix of the true (unobserved) algal abundance in year *t*, **C*_t_*** the matrix of the covariates tested, ice phenology, stratification timing, surface water temperature, and total phosphorus, in year *t*, and **y***_t_* the matrix of the observed chlorophyl *a* concentration in year *t*. Data exploration showed two dominant peaks in chlorophyl *a* concentration in spring and summer/autumn at most sampling stations. To account for spatial and temporal variation in limnological conditions, we evaluated the effects of lake phenology and stratification on algal abundances in spring and summer/autumn separately in three areas of the lake (as above).

### Shifting ice phenology and human disturbances

Lake Simcoe has experienced a host of perturbations induced by human activities over centuries (Evans et al. 1996, North et al. 2013). Excessive nutrient (primarily phosphorous) loading and fishing have been ongoing for more than a century. In the past few decades the onset of shifts in lake ice phenology was also detected (Fig. 1c). To explore how shifting ice phenology propagate through species interactions to shape fish community dynamics, we analyzed the time series data in two steps to evaluate interactive effects of 1) ice phenology and long-term human activities, and 2) ice phenology and limnological properties mediated through plankton communities.

First, we estimated species interactions strengths (**B** in eq. 1) in the fish community using the MARSS model with the selected variance-covariance structures of Q and R above. In **B** the diagonal elements indicate intraspecific interactions (density dependence) strengths, whereas the off-diagonal elements indicate interspecific interactions strengths (Holmes et al. 2023). We used a two-step approach to the **B** matrix estimation. First, prior to model-fitting we selected ecologically plausible species interactions (predation and exploitative competition, (Ives et al. 1999) identified in past research to avoid overparameterization, which were based on field observations of the diets of Lake Simcoe fishes, supplemented by information from the literature (Goto et al. 2020). Because the primary objective of the analyses was to identify the ecological consequences of ice phenology, the selection of species interactions was focused on interactions among five cold-water (numerically dominant) species (lake trout, lake whitefish, cisco, rainbow smelt, and burbot) and their interactions with cool- and warm-water species (yellow perch, northern pike, smallmouth bass, largemouth bass, pumpkinseed, and rock bass, Table 1), resulting in 42 combinations in total. Following the method by (Ives et al. 1999) we evaluated species interaction strengths by first generating 100 sets of randomly generated interactions and then fitting these models to the data. We repeated these processes 100 times (10000 models in total) and identified the most parsimonious model structure using AICc, followed by estimation of confidence intervals using the Hessian approximation (Holmes et al. 2023).

Using the models developed above we analyzed 65-year (1951–2015) time series data of ice phenology and fish community data along with fishing pressure (summer and winter angler effort) and phosphorus loading (mt) with and without species interactions as

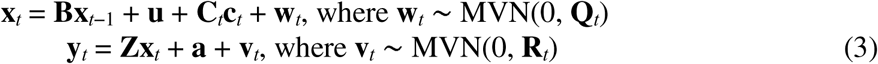

where **x***_t_*, **w***_t_*, and **Q** are the same as eq.1; in the model without species interactions **B** is an *n* × *n* identity matrix, whereas in the model with species interactions **B** is an *n* × *n* interaction matrix (non-zeros in off-diagonal elements); **C*_t_*** is the matrix of coefficients of covariate effects, ice phenology, angler effort, or phosphorus loading on fish abundance in year *t*, **c***_t_* is the matrix of the covariates in year *t*; **y***_t_* is an *n* × 1 matrix of the log-transformed observed abundances (species-specific combined CPUEs estimated in eq. 1) in year *t*, **Z** is an *n* × *n* identity matrix, **a** and **u** are set to 0 (all CPUEs are z-scored), and **v***_t_* is the observation error that follows a multivariate normal distribution with mean 0 and variance-covariance matrix **R**. In model selection we tested for covariates sequentially, ice phenology (ice breakup, freeze-up, and duration) first and if supported then fishing pressure or phosphorus loading (models are partially nested).

### Shifting ice phenology and food web dynamics

To explore how shifting ice phenology propagates through food webs, we analyzed 36-year (1980–2015) time series data on fish and zooplankton abundances, chlorophyl *a* concentration, surface water temperature, and Sacchi depth along with ice and thermal stratification phenology data. First, to evaluate bottom-up effects, in addition to fish species interactions we also incorporated interactions between fishes and zooplankton, in which >80 species were aggregated into four genus groups, Calanoids, Cyclopoids, Cladocera, and Veneroida collected from 12 sampling stations in three areas of the lake (as above); the model structure is the same as eq. 1 except that **x***_t_*, **B**, **u**, **w***_t_*, **y***_t_*, **Z**, **a**, and **v***_t_* matrices now include the zooplankton groups. The **B** matrix was re-estimated using the same two-step approach except that for computation cost and interpretability, only bottom-up interactions from zooplankton to pelagic fishes (including species with planktivorous (larval and juvenile) life stages) were tested (72 combinations in total). And the **C*_t_*** matrix included indirect effects of the covariates through the zooplankton groups. To account for differences in fish spawning season (Table 1) and test for effects of ice and stratification phenologies we averaged zooplankton abundance and limnological covariates over two periods: post-ice breakup spring (April–June) and thermally stratified summer and autumn (August–November). All the covariates were lagged one year to account for surveyed fish communities representing with age 1+ fish. In model selection we tested for covariates sequentially (as above), ice and stratification phenologies first and if supported then limnological properties. All numeric covariates were standardized by subtracting the mean and dividing by the standard deviation (*z*-scores). In species interactions and covariate effects we made inference based only on B and C matrix elements with confidence intervals that do not include zero.

## RESULTS

### Climate effects on shifting ice phenology and thermal profiles

Ice phenology in Lake Simcoe remained stable for the first eight decades of the 20th century (Fig. 1c). Since the early 1980s, however, the timing of ice breakup advanced by 1.7 d decade^-1^ and the timing of ice freeze-up was delayed by 2.6 d decade^-1^, shortening ice cover period by 3.4 d decade^-1^ (Fig. 1c). The most supported model showed that ice phenology was correlated with local daily mean air temperatures in the spring and autumn (prior to ice breakup and freeze-up), which rose by 0.01 and 0.20°C decade^-1^ since the early 1980s (Fig. 1b), but not with any of the regional climate signals tested (Table 2). Shortening ice duration was associated with warmer autumn air temperatures (Fig. 2a).

**Table 2.** Results of model selection for covariates of ice phenology and limnological parameters as response variables in Lake Simcoe. The models in bold indicate the best model for each response variable. AICb and ΔAICb indicate bootstrapped Akaike information criterion scores and differences in AICb scores from the best model.

| response variable | covariate | AICb | ΔAICb | log-likelihood |
| --- | --- | --- | --- | --- |
| ice phenology | base | -34.22 | 19.26 | 29.59 |
|  | spring air temperature | -51.43 | 2.05 | 41.46 |
|  | autumn air temperature | -38.24 | 15.24 | 34.86 |
|  | AMO lag 1 | -30.58 | 22.89 | 31.03 |
|  | AMO lag 2 | -25.38 | 28.10 | 28.43 |
|  | AMO lag 3 | -30.28 | 23.20 | 30.88 |
|  | NAO lag 1 | -29.40 | 24.08 | 30.44 |
|  | NAO lag 2 | -30.03 | 23.45 | 30.76 |
|  | NAO lag 3 | -31.00 | 22.47 | 31.25 |
|  | **spring + autumn air temperature** | **-53.48** | **0.00** | **45.31** |
| stratification phenology | base | -960.2 | 34.2 | 509.9 |
|  | ice breakup | -996.5 | -2.1 | 531.8 |
|  | **post-ice breakup surface water temperature** | **-994.4** | **0.0** | **534.2** |
|  | post-ice breakup bottom water temperature | -821.1 | 173.3 | 491.1 |
| spring chlorophyl *a* (by area) | base | 303.1 | 52.7 | -75.0 |
|  | ice breakup | 255.9 | 5.5 | -36.1 |
|  | **ice breakup + surface water temperature** | **250.4** | **0.0** | **-18.2** |
|  | ice breakup + total phosphorus concentration | 344.5 | 94.0 | -65.3 |
| autumn chlorophyl *a* (by area) | base | 141.5 | 85.4 | 7.0 |
|  | ice breakup | 213.3 | 157.2 | -14.9 |
|  | stratification onset | 145.3 | 89.2 | 20.0 |
|  | stratification duration | 108.0 | 51.9 | 38.7 |
|  | **stratification turnover** | **56.1** | **0.0** | **64.6** |
|  | stratification duration + surface water temperature | 233.1 | 176.9 | -8.4 |
|  | stratification turnover + surface water temperature | 168.2 | 112.1 | 24.1 |

**Figure 2.**
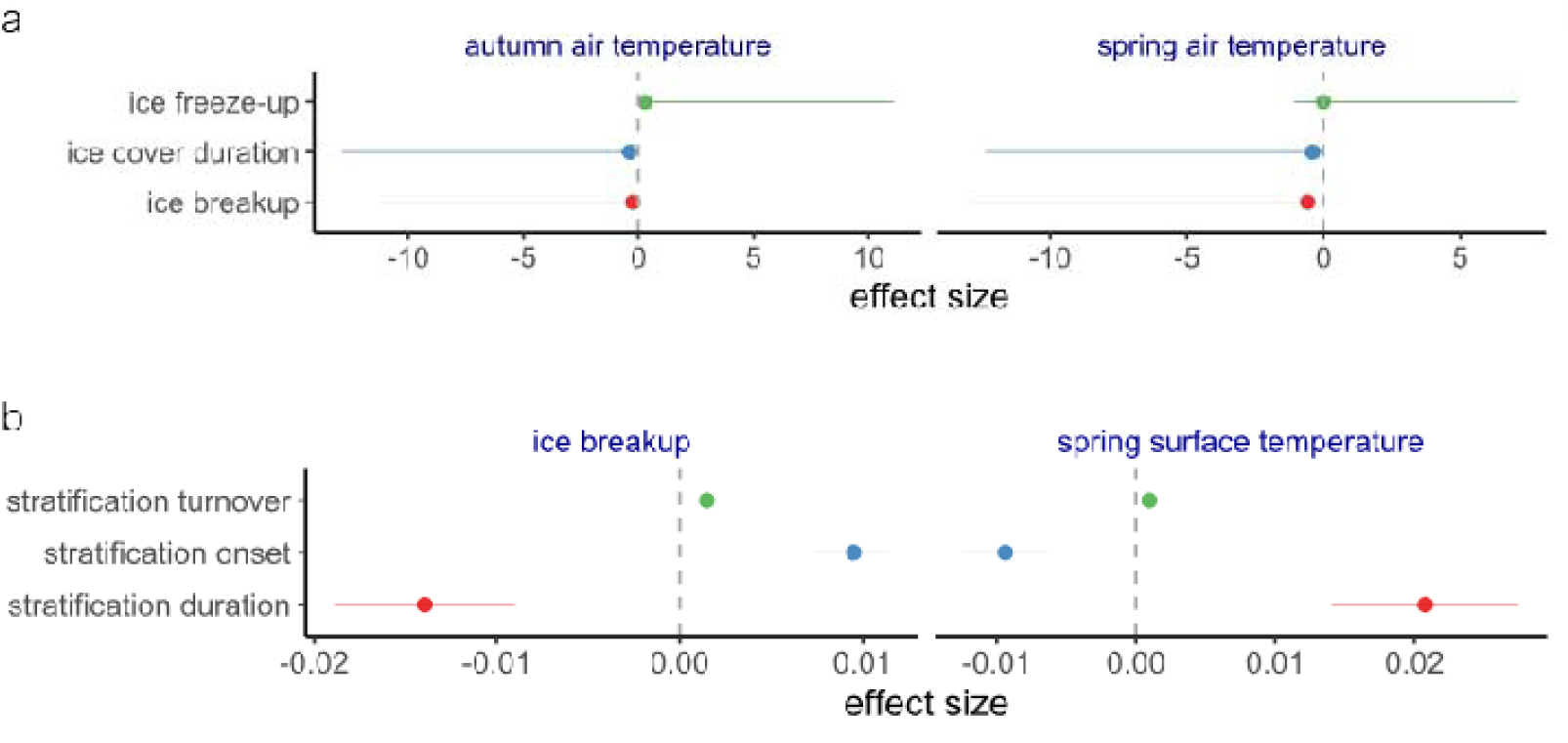
Drivers of ice phenology and thermal stratification phenology in Lake Simcoe. (a) Climate (pre-ice breakup air temperature) effects on ice phenology (ice cover duration, ice breakup, and ice freeze-up) and (b) ice breakup timing and air temperature effects on thermal stratification phenology (duration, onset, and turnover). Effect sizes are estimated using MARSS (multivariate autoregressive state-space) models applied to ice and thermal stratification time series data.

The timing of stratification onset advanced by on average 4.1, 3.2, and 4.0 d decade^-1^ in nearshore, offshore, and Kempenfelt Bay and the timing of turnover was delayed by 5.9, 3.3, and 2.8 d decade^-1^, prolonging stratification period by 9.7, 6.5, and 6.3 d decade^-1^ between 1980 and 2012 (Fig. 1d). The most supported model showed that the timing of stratification advanced with that of earlier ice breakup and with warmer surface water temperatures in spring (Fig. 2b and Table 2).

### Effects of phenological and limnological changes on seasonal algal blooms

The best model showed that advancing timings of lake ice breakup contributed to declining spring algal abundances (estimated by chlorophyl *a* concentration) in offshore and Kempenfelt Bay (Fig. 3a and Table 2). Likewise, declining spring algal abundances were correlated with warming surface water temperatures in all areas (Fig. 3a), whereas declining total phosphorus concentration contributed little after accounting for early ice breakup (Table 2). By contrast, shifting ice phenology contributed little to autumn algal abundances (Table 2). Delayed lake stratification turnover (in turn longer stratification period) was correlated with declining algal abundances in nearshore but rising algal abundances in offshore and Kempenfelt Bay (Fig. 3b). Warming surface water temperatures contributed little to declining algal abundances after accounting for stratification phenology changes (Table 2).

**Figure 3.**
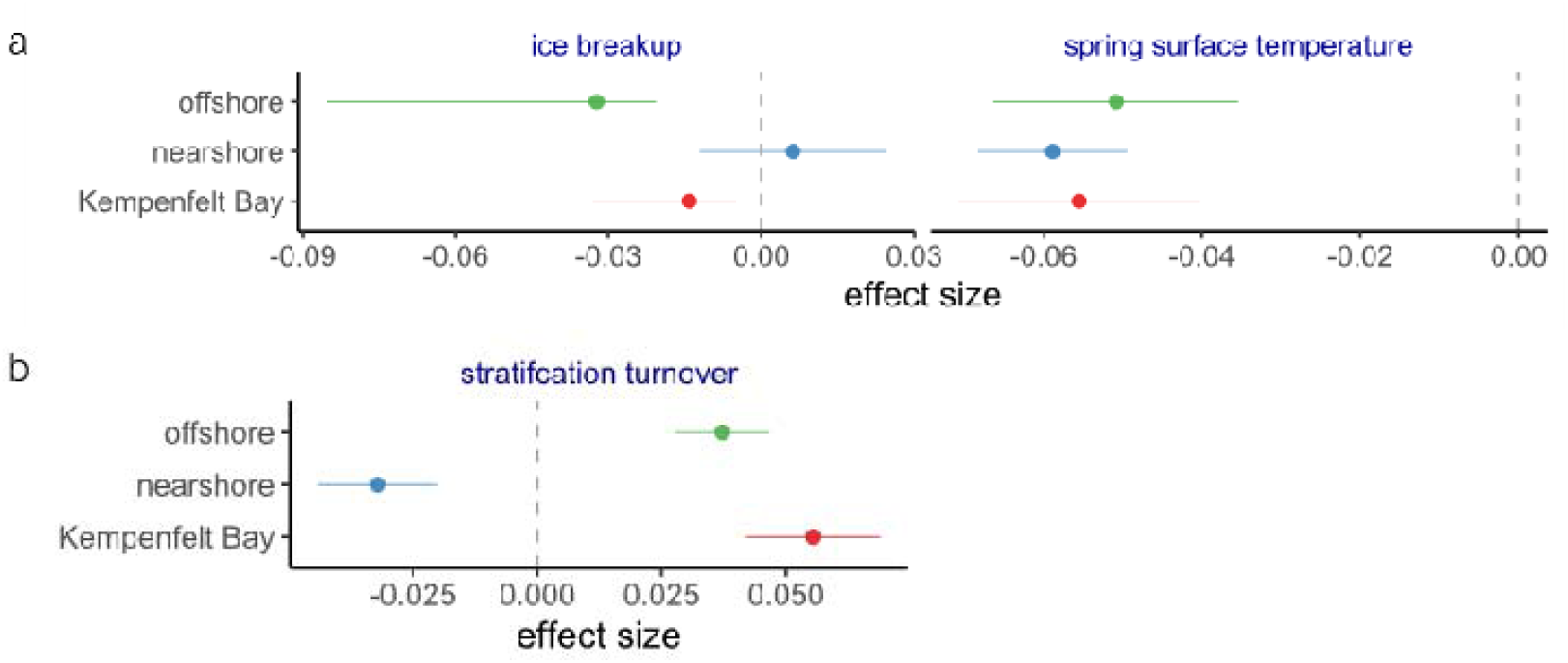
Drivers of spring and autumn algal production in Lake Simcoe. (a) Ice breakup timing and spring (post-ice breakup) surface water temperature effects on spring chlorophyl *a* concentration in concentration in Kempenfelt Bay and offshore and nearshore areas and (b) thermal stratification turnover timing effect on autumn chlorophyl *a* concentration in concentration in Kempenfelt Bay and offshore and nearshore areas. Effect sizes are estimated using MARSS (multivariate autoregressive state-space) models applied to chlorophyl *a* concentration time series data.

### Reconstruction of fish community dynamics

Reconstructed fish community dynamics in Lake Simcoe showed that most species had experienced big relative changes (CV = 0.46 to 1.46) in abundance over 65 years (Fig. 4a), based on the most supported model (with unconstrained and species-specific variance–covariance structures in state and observation, Table S1). The population sizes of intensely harvested species like lake trout and lake whitefish declined on average more than 80% from the 1960s to the 2000s before starting to recover (Fig. 4a). During the same period the population sizes of other top predators (e.g., largemouth bass) and mesopredators (e.g., yellow perch) rose as much as 8.1-fold (Fig. 4a). The best model showed that the process variance of the fish community dynamics varied among species (σ^2^ = 0.004 to σ 0.52) and were correlated between years; the observation variance also varied among species and surveys (σ^2^ = 0.12 to 0.26).

**Figure 4.**
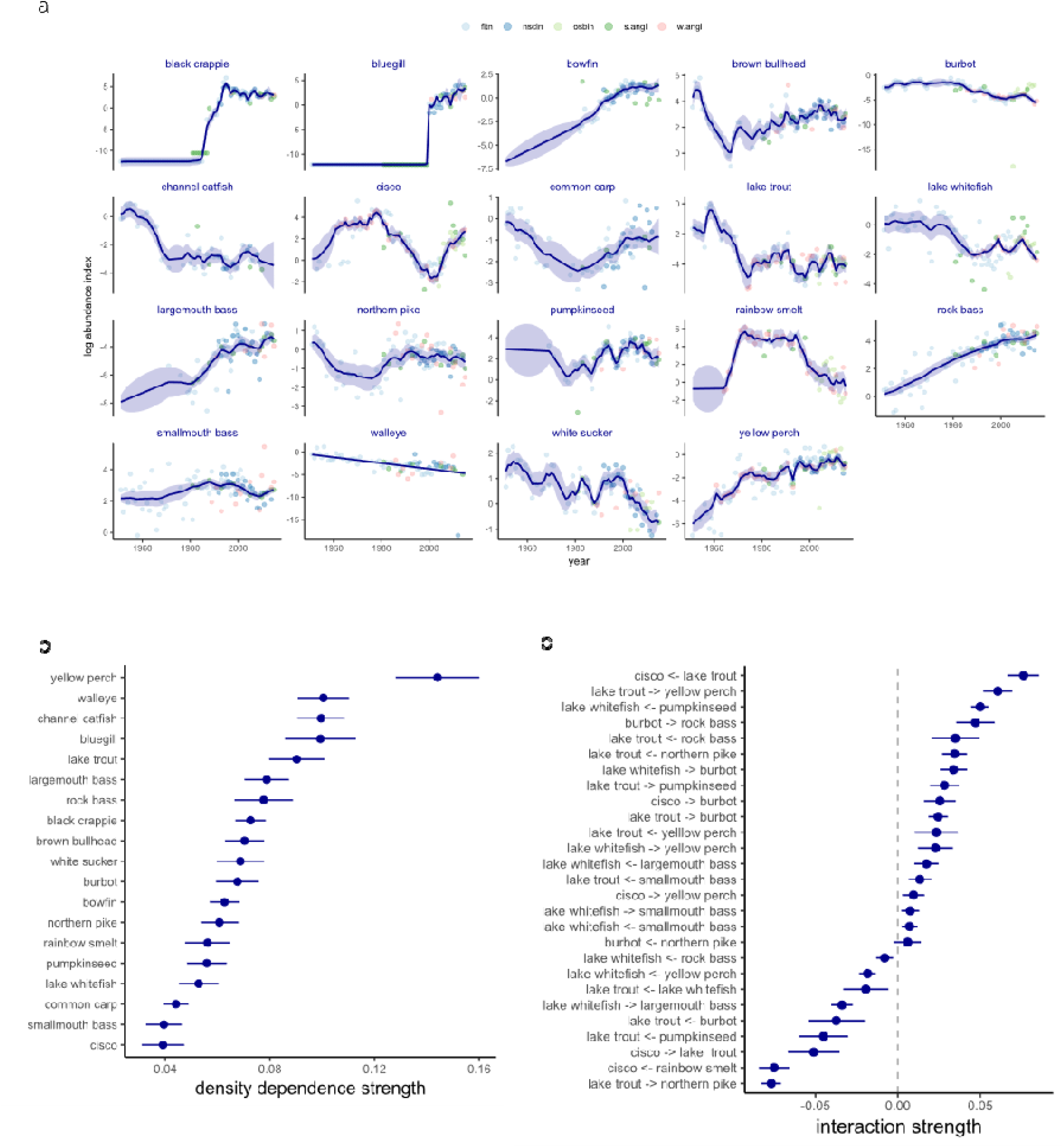
Reconstructed Lake Simcoe fish community dynamics in 1951–2015 using a MARSS (multivariate autoregressive state-space) model. (a) Scaled log-transformed abundance index of 19 species (black crappie, bluegill, bowfin, brown bullhead, burbot, channel catfish, cisco, common carp, lake trout, lake whitefish, largemouth bass, northern pike, pumpkinseed, rainbow smelt, rock bass, smallmouth bass, walleye, white sucker, and yellow perch), (b) density dependence strengths of the 19 species, and (c) interaction strengths of select species pairs. The model was fitted to five fishery–independent and –dependent time series data (FITN, OSBIN, NSCIN, and winter and summer fisheries CPUEs).

### Effects of shifting ice phenology and human disturbances on fish community dynamics

The most supported fish community model comprised 27 interactions, of which 24 were supported by the data (confidence intervals did not include zero, Fig. 4c). These interactions showed numerically dominant cold-water species like lake trout and lake whitefish negatively affected cool- and warm-water northern pike and largemouth bass, whereas cool- and warm-water mesopredator species like yellow perch and pumpkinseed negative affected the cold-water species (Fig. 4c). The model also showed a negative effect of rainbow smelt (an invasive cold-water planktivore) on cold-water cisco, indicating competition for shared resources. Some interactions like positive effects on competing species or predators on prey suggest indirect effects (predators feeding on competing species of prey for example). Lake Simcoe fish species were in general weakly (1 – B matrix diagonal parameters < 0.1) regulated by density dependence, with yellow perch being relatively more density-regulated (Fig. 4b).

Shifting ice phenology consistently contributed to the population dynamics of most fish species; contrary effects of advancing timings of ice breakup and delaying timings of ice freeze-up on most species with varying thermal preferences (Fig. 5 and Table 3). For example, earlier ice breakups were correlated negatively with cold-water lake trout and several warm-water population growth but positively with cold-water burbot (Fig. 5a). In contrast, later ice freeze-ups were correlated positively with cold-water lake trout and cisco population growth but negatively with lake whitefish (Fig. 5b,c). Likewise, later ice freeze-ups were correlated positively with warm-water largemouth bass but negatively with northern pike and cold-water burbot (Fig. 5b,c). With the ice freeze-up effects accounted for, fishing effort by anglers in both winter and summer was correlated negatively with cold-water species population growth and positively with many warm-water species population growth (Fig. 5b,c). The models without species interactions accounted for showed consistent patterns in the directions of covariate effects with those with species interactions but in general overestimated the strengths of covariate effects.

**Figure 5.**
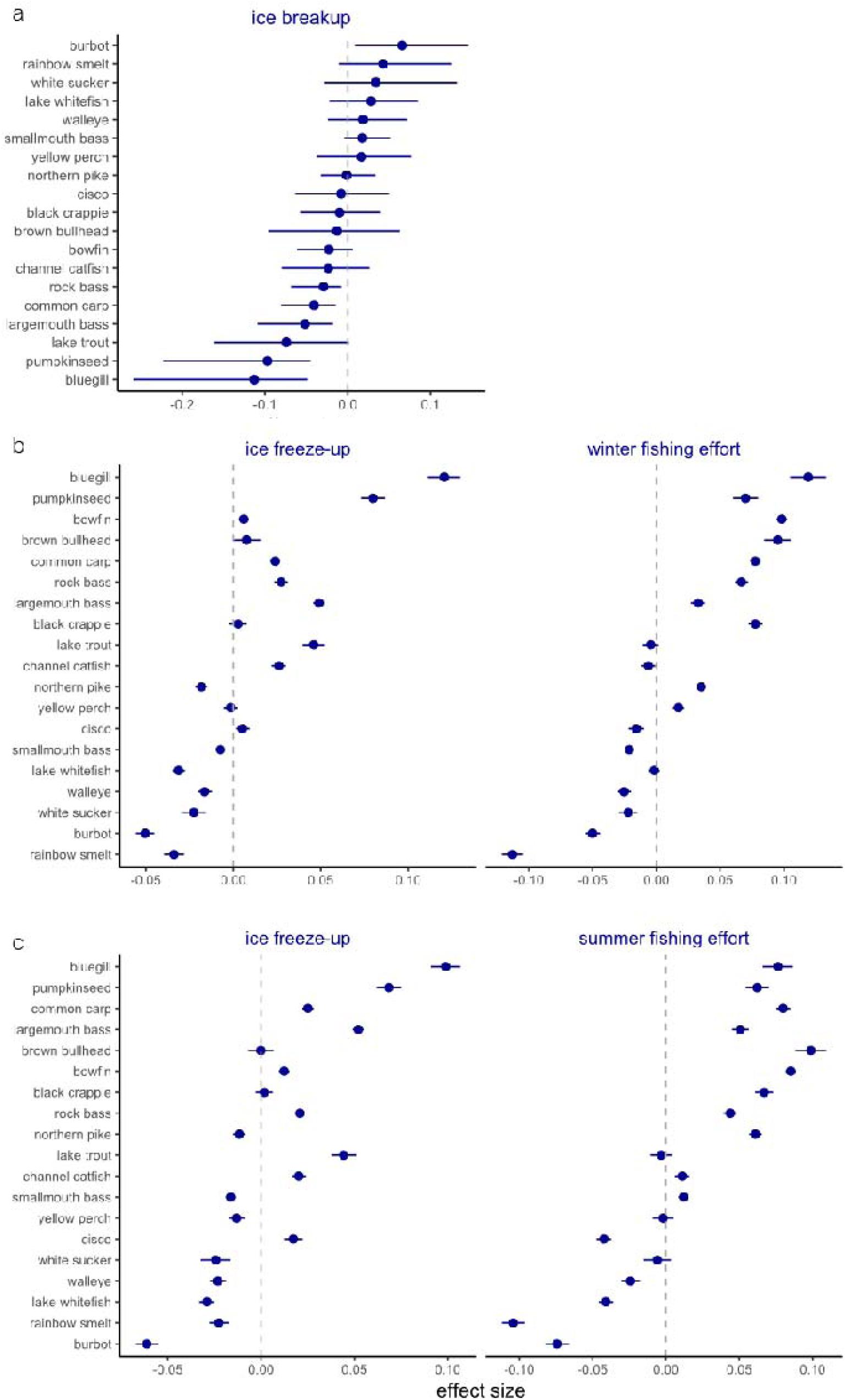
Drivers of fish community dynamics in Lake Simcoe. (a) Ice freeze-up timing and winter fishing effort (angler hours) effects on the population growth of the 19 species and (b) ice freeze-up timing and summer fishing effort (angler hours) effects on the population growth of the 19 species. Effect sizes are estimated using MARSS (multivariate autoregressive state-space) models applied to fish relative abundance (CPUE) time series data.

**Table 3.** Results of model selection for covariates of fish community as response variable in Lake Simcoe. AICb indicate bootstrapped Akaike information criterion scores. The models are partially nested because they are evaluated to test for only ecologically meaningful combinations of ice phenology and fishing pressure or phosphorus loading.

| covariate | AICb | log-likelihood |
| --- | --- | --- |
| base | -934.8 | 819.9 |
| ice duration | -833.8 | 801.4 |
| ice breakup | -1823.4 | 1296.2 |
| ice freeze-up | -1088.3 | 928.7 |
| ice breakup + phosphorus loading | -1286.1 | 1061.0 |
| ice breakup + winter angling | -1299.7 | 1067.8 |
| ice breakup + summer angling | -1295.0 | 1065.4 |
| ice freeze-up + winter angling | -1181.6 | 1008.7 |
| ice freeze-up + summer angling | -1417.2 | 1126.5 |

### Bottom-up effects of shifting ice phenology via food web dynamics

Reconstructed food web dynamics in Lake Simcoe most supported by the model (Table S2) showed that most zooplankton groups declined by 46–73% except invasive Veneroida (primarily dreissenid vilegers), whose abundances rose 3.2–6.2 fold (Fig. 6a). The zooplankton groups were weakly (1 – B matrix diagonal parameters = 0.13–0.25) regulated by density dependence, except Veneroida, which was more strongly (0.33–0.96) regulated by density dependence (Fig. 6b). The best model comprised 41 additional fish–zooplankton interactions, of which six were statistically supported (Fig. 6c). These interactions highlighted an increasingly dominant role of Veneroida in the food web, which promoted some cool- and warm-water fish population (especially yellow perch and largemouth bass) growth and attenuated cold-water cisco growth (Fig. 6c).

**Figure 6.**
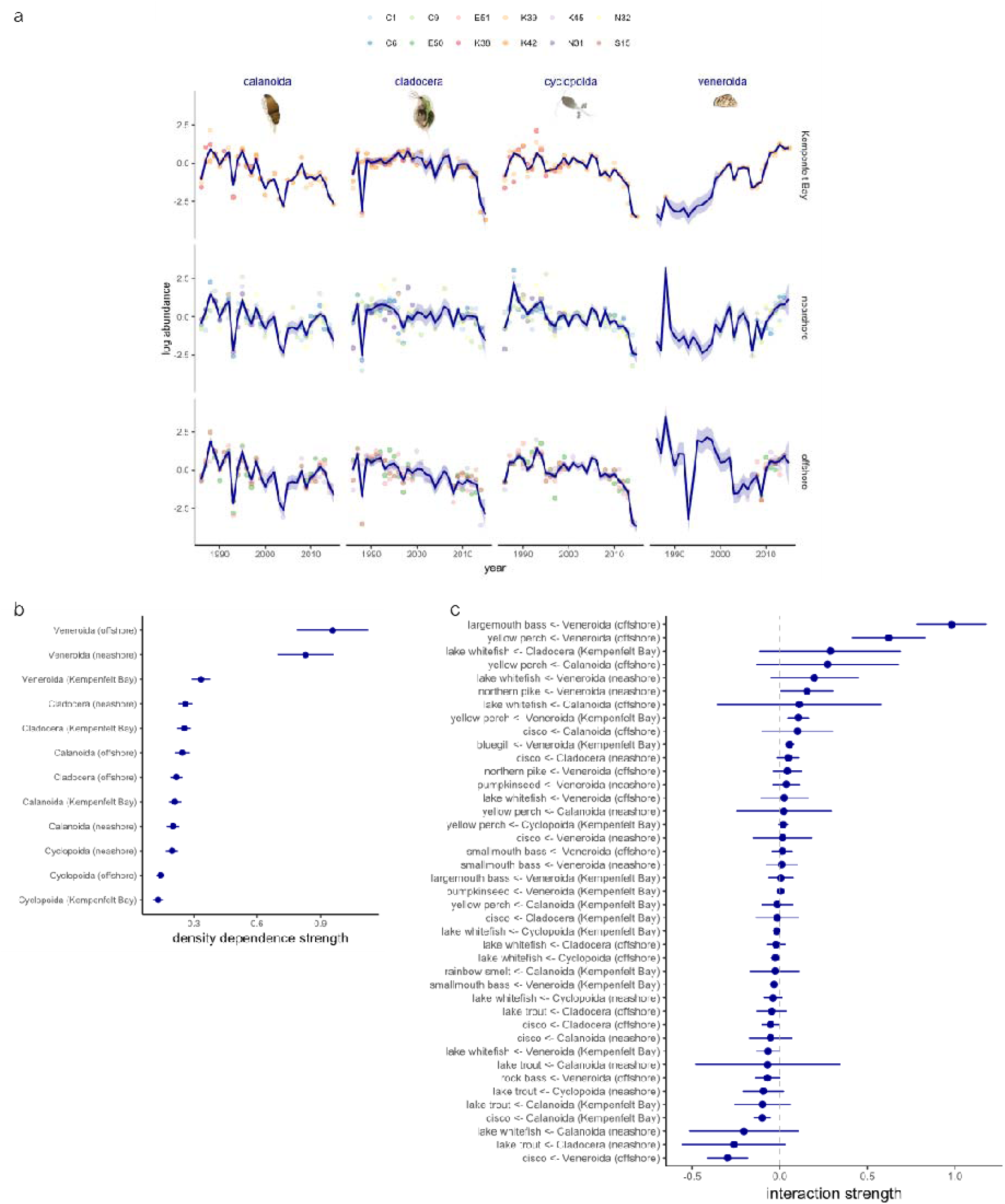
Reconstructed Lake Simcoe food web dynamics during 1980–2015 using a MARSS (multivariate autoregressive state-space) model. (a) Scaled log-transformed abundance index of four major zooplankton groups (Calanoids, Cyclopoids, Cladocera, and Veneroida) in Kempenfelt Bay and offshore and nearshore areas and (b) density dependence strengths of the four zooplankton groups in the three areas, and (c) interaction strengths between select fish and zooplankton groups. The model was fitted to fish relative abundance (CPUE) and zooplankton abundance time series data.

Advancing ice breakup timings were negatively correlated with the growth rates of most zooplankton groups lake-wide with a few exceptions in deeper Kempenfelt Bay, where Cladocera and Veneroida growth rates were positively correlated with earlier ice breakups (Fig. 7a, Table 4). Prolonging stratification periods negatively affected most zooplankton groups especially in nearshore areas (Fig. 7b). After accounting for ice phenology, warming surface water temperatures in the spring promoted the growth rates of many zooplankton groups (Fig. 7a). After accounting for stratification phenology, deepening Secchi depth was more supported by data than declining algal abundances (Table 4), reducing most zooplankton (except Veneroida in offshore and Kempenfelt Bay) growths (Fig. 7b).

**Figure 7.**
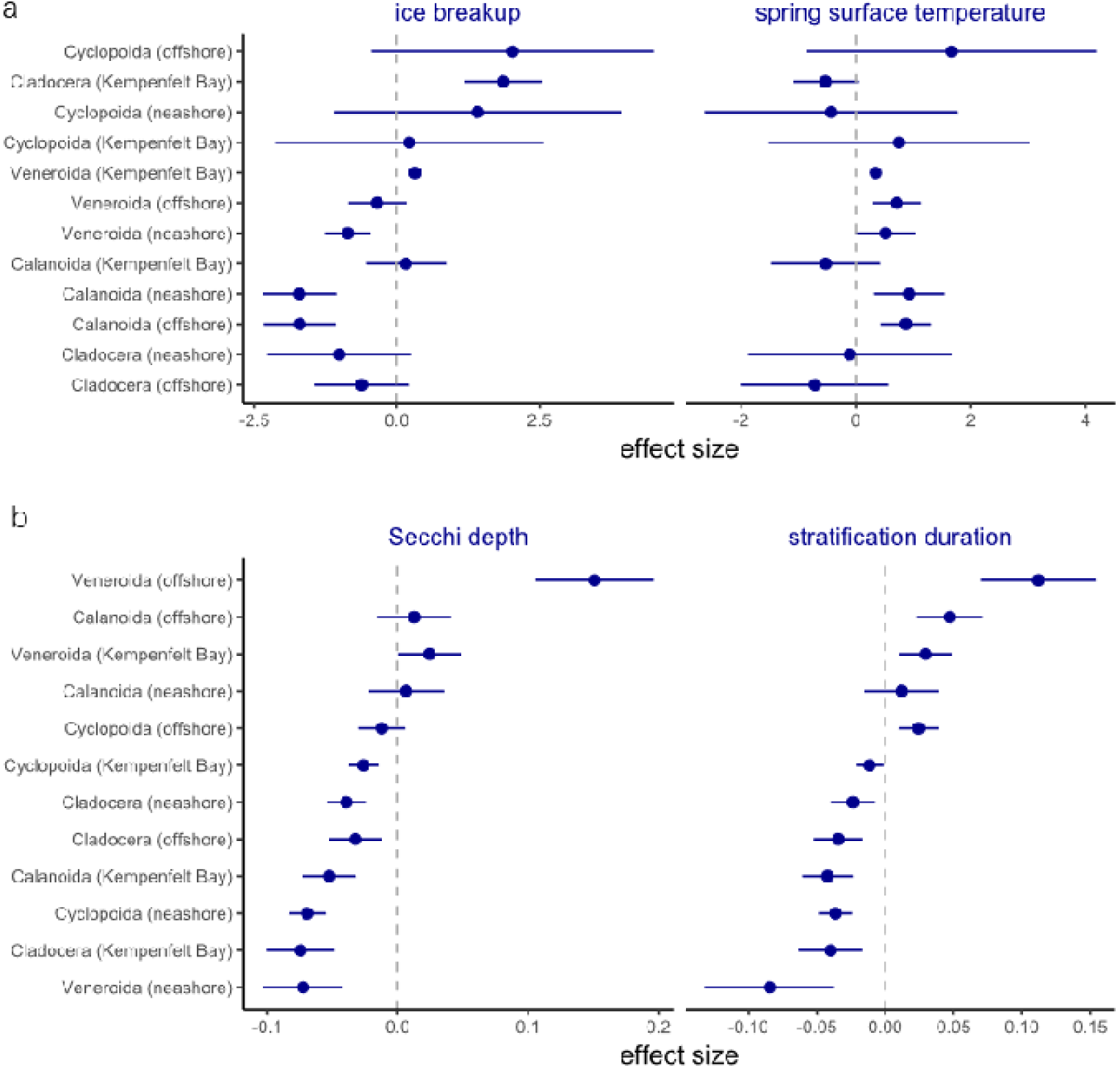
Drivers of zooplankton community dynamics in the Lake Simcoe food web during 1980–2015. (a) Ice breakup timing and spring surface water temperature on the population growth of four zooplankton groups (Calanoids, Cyclopoids, Cladocera, and Veneroida) in Kempenfelt Bay and offshore and nearshore areas and (b) water clarity (Secchi depth) and thermal stratification timing on the population growth of four zooplankton groups in the three areas. Effect sizes are estimated using MARSS (multivariate autoregressive state-space) models applied to fish relative abundance (CPUE) and zooplankton abundance time series data.

**Table 4.** Results of model selection for covariates of food web dynamics as response variable in Lake Simcoe. AICb indicate bootstrapped Akaike information criterion scores. The models are partially nested because they are evaluated to test for only ecologically meaningful combinations of ice or stratification phenology and limnological parameters.

| covariate | AICb | log-likelihood |
| --- | --- | --- |
| base | 889.0 | -159.0 |
| ice duration | 1107.8 | -248.4 |
| ice freeze-up | 727.8 | -58.4 |
| ice breakup | 602.0 | 4.5 |
| stratification duration | 847.8 | -118.4 |
| stratification onset | 985.5 | -187.3 |
| stratification turnover | 1023.5 | -206.3 |
| ice breakup + surface water temperature | 563.7 | 44.2 |
| ice breakup + chlorophyl *a* concentration | 909.0 | -128.5 |
| stratification duration + surface water temperature | 1137.7 | -242.8 |
| stratification duration + chlorophyl *a* concentration | 810.1 | -79.0 |
| stratification duration + Secchi depth | 750.9 | -49.4 |

## DISCUSSION

Our analysis integrating over 140 historical and contemporary time series of climate, physical, and biological data revealed that climate change-induced shifts in ice phenology in recent decades can trigger limnological and ecological changes that propagate through pelagic food webs, reshaping fish community and fisheries dynamics in northern temperate lakes. Large, relatively shallow lakes like Lake Simcoe are especially prone to climate change-induced ice loss (Woolway et al. 2020). Increasingly earlier ice breakups, later ice freeze-ups, earlier thermal stratification onset, and later turnover since the early 1980s, along warming water temperatures, attenuated algal production. Although the declines in algal abundance to some extent contributed to declining zooplankton population growth, shifting ice phenology, combined with increasingly more abundant invasive dreissenids, likely played larger roles in most zooplankton groups’ population dynamics. Increasingly greater water clarity (less pelagic algae), along with longer stratification duration, likely exposed the zooplankton to greater predation pressure.

Shifting ice phenology further modified the population and community dynamics of fish species with differential thermal preferences, interactively with time-varying non-climate drivers. Advanced timings of ice breakup and delayed timings of ice freeze-up promoted and diminished their population growth. Although management interventions–reductions in nutrient loading and fishing effort in particular–in recent decades aided cold-water fish species recoveries, their interaction patterns and strengths with cool- and warm-water species were further modulated by phenological shifts in limnological condition filtered through plankton communities. These findings reveal how complex food web dynamics can emerge from ecologically divergent responses that reshape species interactions within and among trophic levels under shifting ice phenology filtered through bottom-up processes, likely buffering destabilizing effects of a changing climate.

### Lake ice phenology and population dynamics

Timings of lake ice breakup often coincide with seasonal life history events of many aquatic species (Woods et al. 2022). Climate warming-induced shifts in timing of spring ice breakup can trigger a series of cascading changes in physical and in turn biological (life history and demographic) processes that regulate the population dynamics of many species in northern temperate lakes (Straile et al. 2003, Winder and Schindler 2004b, Mishra et al. 2011). In Lake Simcoe increasingly earlier ice breakups and warmer surface water temperatures under recent decades’ climate warming coincided with reductions in spring algal production especially in offshore waters. As the timings of spring ice breakup and subsequent water temperature warming advanced since the 1980s, primary producers in many temperate lakes also experienced shifts in their reproductive phenology, as also documented for Lake Washinton (USA, Winder and Schindler 2004b), Lake Windermere (UK, George and Taylor 1995), and Lake Baikal (Russia, Hampton et al. 2008). Because winter ice cover reduces light penetration, inhibiting algal growth, ice phenology can also regulate the phenology of algal growth (Straile et al. 2003, Woolway et al. 2021b). Climate change could thus enhance the role of environment drivers in, and in turn amplify variability and reduce predictability in population dynamics of these species with seasonal life histories (Post 2013, 2019).

Warming climate-induced earlier ice breakup and surface water warming can also trigger earlier onset of thermal stratification (Winder and Schindler 2004b). Warmer air and surface water temperatures further strengthen the stratification prolonging its period (Winder and Schindler 2004b, Woolway et al. 2021b) and in turn delaying freeze-up timing (Li et al. 2022). In Lake Simcoe prolonged stratification periods, combined with attenuated algal production with increasingly earlier peaks to some extent, contributed to reduced abundances of zooplankton especially in nearshore areas, likely owing to higher metabolic cost induced by warmer temperatures (Straile et al. 2003). Intensified stratification in lakes can reduce the frequency of mixing (Woolway and Merchant 2019) and disrupt nutrient cycling, diminishing lake productivity (O’Reilly et al. 2003, Cohen et al. 2016).

Advancing ice breakup and delaying freeze-up timings would prolong growing seasons in northern temperate lakes, resulting in varying consequences among fish species with differential thermal preferences (Helland et al. 2011, Shuter et al. 2012, McMeans et al. 2020), including changes in reproduction phenology (Shuter et al. 2012, Farmer et al. 2015, Hovel et al. 2017) and life history traits (like migration timing, Cline et al. 2019). In Lake Simcoe weak density dependency in fish populations suggesting high sensitivity to extrinsic drivers like shifting ice phenology. Increasingly earlier ice breakups in spring promoted the population growth of some fishes (including lake whitefish and smallmouth bass). Earlier ice breakups can advance spawning timings (Hovel et al. 2017, Tao et al. 2018, Feiner et al. 2022), allowing larval and juvenile fish to grow more over longer periods if their food needs are also met (Schindler et al. 2005, Hovel et al. 2017) and thus enhancing their survival probabilities later (Feiner et al. 2022). Shortening ice cover periods may also attenuate the severity of winter kills–a mass mortality event caused by oxygen depletion in northern temperate lakes (Shuter et al. 2012, McMeans et al. 2020). Shorter winter may, however, induce adverse effects on reproductive biology for some species, including lower egg viability and larval survival (Farmer et al. 2015).

Longer growing seasons resulting from shifting ice phenology may not necessarily benefit all species; warmer water temperatures during ice-free seasons can also alter seasonal habitat use for fish, promoting or attenuating population growth depending on thermal preferences (Tunney et al. 2014). Warming for example can create stressful conditions in littoral habitat for cold-water fish like salmonids, forcing them to remain offshore (Guzzo et al. 2017, Caldwell et al. 2020, McMeans et al. 2020). As productive littoral habitat becomes less suitable for cold-water fish under climate warming, these fish may resort to remain offshore, relying more on pelagic resources (Caldwell et al. 2020). In Lake Simcoe earlier ice breakups adversely affected the population growth of several fish species including largemouth bass and lake trout. These fishes may have experienced unfavorable habitat conditions; for example, advanced ice breakup dates can have detrimental effects on young fish (Feiner et al. 2022) and in turn population growth if food needs were not met. Longer and stronger thermal stratification duration can also amplify the severity of oxygen depletion in bottom waters, preventing fishes from accessing their food supply (Shuter et al. 2012, Goto et al. 2017). In Lake Simcoe the fishes that benefited from earlier ice breakups (burbot for example) experienced the opposite from later ice freeze-ups in the autumn.

### Life history event timings and species interactions

Shrinking ice cover and expanding stratification periods as climate warms may fundamentally restructure lake food webs (Shuter et al. 2012, McMeans et al. 2020). Advanced timings of spring algal blooms, for example, can support higher metabolism in zooplankton triggered by surface water warming (Straile et al. 2003). In Lake Simcoe spring zooplankton growth reflected declining algal production both responding to increasingly earlier ice breakups. The algal–grazer relationships, however, became weakened in the summer. The effects of shifting lake phenology can be attenuated as it moves up the food chain (Thackeray et al. 2013, Thackeray et al. 2016, Cohen et al. 2018), likely resulting from differential responses to the shift. Differential shifts in phenology among trophic levels can disrupt or even decouple trophic interactions (‘trophic mismatch’, Winder and Schindler 2004b, de Senerpont Domis et al. 2007, Jan et al. 2024). In Lake Simcoe, however, effects of shifting ice phenology may have played out over the ice-free season through limnological processes and trophic interactions jointly. Warmer post-ice breakup water temperatures and prolonged stratification durations in the summer (and early autumn) promoted the population growth of some zooplankton groups (including Veneroida and Calanoida in offshore waters) and in turn fish species with all thermal preferences. Stronger (by an order of magnitude) bottom-up effects of increasingly more abundant dreissenid veligers (planktonic larvae) in particular enhanced the population growth of fishes like yellow perch and lake whitefish lake-wide. Because adults of these species consume adult dreissenids in the lake (Goto et al. 2020), these relationships may also reflect their increasing reliance on dreissenids under a warming climate. For example, earlier ice breakups, warmer water temperatures, and longer stratification durations collectively aided the population growth of dreissenid veligers and in turn yellow perch.

Effects of ice phenology may also be context-dependent; species may respond only when certain physical or ecological conditions are present (Helland et al. 2011, McMeans et al. 2020). In Lake Simcoe interaction strengths among zooplankton species, which have limited horizontal movement (more localized species interactions), were relatively high; invasive dreissenid veligers in particular show increasingly stronger intra- and inter-specific interactions as their abundance ballooned following the invasions in 1993 (McNeice and Johanson 2004). Although delayed timings of ice freeze-up (longer growing seasons) promoted the growth of some zooplankton groups like cyclopoid copepods, greater water clarity owing to improved water quality through reductions in phosphorus loading, combined with high filtering rates of adult dreissenids depleting pelagic algae (Hecky et al. 2004), may have exposed zooplankton to higher predation pressure (Bunnell et al. 2021) over longer stratification periods, further contributing to their recent population declines. By contrast, consistently negative effects by dreissenid vilegers on planktivorous cisco likely indicate indirect effects mediated by adult dreissenids filtering algae, depleting food supply for other zooplankton groups.

Climate change may also promote synchrony in phenology within a trophic level (Post 2013, Kharouba et al. 2018, Post 2019) as opposed to asynchrony among trophic levels (Thackeray et al. 2013). Although some cold-adapted species like lake trout in the Arctic and subarctic may benefit from expanded habitats with less competition under warming climates (Campana et al. 2020), shortening of ice cover periods in northern temperate lakes may promote synchrony in seasonal habitat use, amplifying competition for shared resources (Helland et al. 2011, Guzzo et al. 2017, McMeans et al. 2020). In Lake Simcoe our analysis reveals divergent responses to advanced timings of ice breakup and delayed timings of freeze-up among fish species and consequences for species interactions. Negative species interactions detected in our analysis, for example, are partially explained by their differential responses to earlier ice breakups; species that were adversely affected by earlier ice breakups (like lake trout and largemouth bass) experienced density-dependent effects from species that benefited from earlier ice breakups (like lake whitefish and burbot). Seasonal timings of life history events in many species have developed to optimize fitness for persistence and coexistence (Post 2013, White and Hastings 2020), through promoting synchrony (overlap between consumers and resources, Post 2013, 2019) or asynchrony (avoidance of competition for shared resources, Helland et al. 2011, Post 2019, McMeans et al. 2020). Shifting ice phenology in north temperate lakes can, for example, trigger time allocation shifts in life history events; shorter spawning and larval development periods for cold-water fishes and longer spawning and development periods for cool- and warm-water fishes (Shuter et al. 2012). Although spatial variation in ice phenology and limnological properties may buffer their effects on fish species to some extent, as shown in our analysis, shorter winters can also intensify competition for shared resources (like spawning grounds) required for life history events (Post 2019, McMeans et al. 2020). Directional changes in these life history events of two or more interacting species may thus reshape food web dynamics and stability (Post 2013, 2019, Zhao et al. 2023) under shifting ice phenology.

### Food web responses to shifting lake phenology modulated by human activities

Like many northern temperate lakes Lake Simcoe has experienced, along with climate warming, perturbations from a host of human activities for centuries, including cultural eutrophication, overharvesting, and species invasion (Evans et al. 1996, North et al. 2013, Dunlop et al. 2019). Despite favorable conditions created by shifting ice phenology under climate warming, unlike other systems (Carter and Schindler 2012), plankton abundances in Lake Simcoe have been declining the past few decades. This pattern partly results from ongoing management-assisted oligotrophication, which was accelerated following the invasion of dreissenids owing to their high abundance and filtration rate (North et al. 2013).

Stricter regulation on nutrient loadings, combined with reduced fishing pressures, also aided the recoveries of Lake Simcoe cold-water fishes like lake trout, lake whitefish, and cisco, highly sought-after species in winter fisheries, after decades of depleted status (Goto et al. 2020). Climate change-induced shifting ice phenology may, however, be hampering the recoveries of these fishes by systematically altering the lake’s physical and ecological environments. Our analysis shows that shifting ice phenology and fishing interactively regulated the population dynamics of fish species with varying thermal preferences differentially, modulating their interactions and food web dynamics. Many Lake Simcoe fish populations experience high fishing pressure annually especially in winter (Dunlop et al. 2019). Shorter ice cover periods can however shift fishing pressure from winter to summer, amplifying or attenuating the strength of species interaction both vertically and horizontally (in addition to greater activities of warm- and cool-water species in the winter, Helland et al. 2011, Guzzo et al. 2017, McMeans et al. 2020). These indirect effects of shifting ice phenology can also propagate through the food web in the winter, perhaps having carry-over effects in the spring. Climate change-induced shifts in fishing pressure may thus directly or indirectly modify the population dynamics of harvested species and their interactions with others while also posing challenges in deciphering causal mechanisms underlying the fish community dynamics under climate warming.

### Managing shared living resources under shifting phenologies

Many northern temperate lakes support a variety of ecosystem services, including fisheries production for recreation, commerce, and subsistence (Welcomme et al. 2010, Lynch et al. 2016). Continued climate change-induced shifts in ice phenology and winter severity will likely modify how extrinsic pressures shape how aquatic populations grow and interact with one another (Dinh et al. 2023), and in turn human activities around lakes (Cline et al. 2019). Although variability in the phenology of ice phenology and thermal stratification has not been detected in Lake Simcoe unlike some northern temperate lakes (Feiner et al. 2022), this may change in the coming decades based on the projections by recent research (Filazzola et al. 2024). Increases in climate change-induced variability in lake phenology in northern temperate lakes would likely ripple through food webs, as our study demonstrated, and further reshape population and community dynamics, altering ecosystem processes and services like winter ice fishing (Filazzola et al. 2024). In Lake Simcoe recreational ice fishing can generate more than US$20 million annually (Filazzola et al. 2024). Projected changes in ice phenology and resulting declines in winter ice cover duration would thus reduce fishing pressure especially cold-water predatory fishes like lake trout, which could further modify species interaction dynamics and top-down control of food webs in the winter, whose effects may be carried over to the spring. Accounting for shifting phenologies in limnological condition, species interaction, and human activity would thus not only help deepen our understanding of food web dynamics in northern temperate lakes under a changing climate but also develop robust policies that safeguard against climate-induced risks and manage fisheries resources sustainably.

## ACKNOWLEDGEMENTS

We thank the Lake Simcoe Fisheries Assessment Unit of the Ontario Ministry of Natural Resources and Forestry (MNRF) and the Ontario Ministry of the Environment, Conservation and Parks (MECP) for field and laboratory work and data management for Lake Simcoe long-term monitoring programs. We thank Justin Trumpickas (MNRF) and Hamdi Jarjanazi (MECP) for compiling and providing the field survey data. Some figures use images from the IAN Symbols, courtesy of the Integration and Application Network, University of Maryland Center for Environmental Science (ian.umces.edu/symbols/).

---

**Table S1.** Results of model selection for model structure of fish community dynamics in Lake Simcoe during 1950–2015. The models in bold indicate the best model for each response variable. **u**, **Q**, and **R** indicate matrices of the long-term mean population growth rates of 19 species, and the variance-covariance of the process error and the observation error with a multivariate normal distribution. AICb and ΔAICb indicate bootstrapped Akaike information criterion scores and differences in AICb scores from the best model.

| model | **u** | **Q** | **R** | AICb | ΔAICb | log-likelihood |
| --- | --- | --- | --- | --- | --- | --- |
| base | equal | diagonal and equal | diagonal and equal | 3991.3 | 344.5 | -1913.7 |
| M1 | unequal | diagonal and equal | diagonal and equal | 4011.7 | 364.8 | -1903.8 |
| M2 | unequal | diagonal and unequal | diagonal and equal | 4017.6 | 370.8 | -1886.2 |
| M3 | unequal | unconstrained | diagonal and equal | 4150.1 | 503.3 | -1757.0 |
| M4 | unequal | diagonal and equal | diagonal and unequal | 3728.5 | 81.7 | -1674.3 |
| M5 | unequal | diagonal and unequal | diagonal and unequal | 3709.7 | 62.9 | -1642.2 |
| M6 | **unequal** | **unconstrained** | **diagonal and unequal** | **3646.9** | 0.0 | **-1462.7** |
| M7 | unequal | diagonal and equal | grouped by species | 3945.5 | 298.7 | -1850.1 |
| M8 | unequal | diagonal and unequal | grouped by species | 3950.4 | 303.5 | -1853.7 |
| M9 | unequal | unconstrained | grouped by species | 4126.8 | 479.9 | -1690.9 |
| M10 | unequal | diagonal and equal | grouped by survey | 3935.5 | 288.6 | -1861.1 |
| M11 | unequal | diagonal and unequal | grouped by survey | 3919.7 | 272.8 | -1832.5 |
| M12 | unequal | unconstrained | grouped by survey | 4022.6 | 375.8 | -1659.8 |

**Table S2.** Results of model selection for model structure of the food web dynamics in Lake Simcoe during 1980–2015. The model structure of the fish community component is estimated above (Table S1) and the parameters are fixed in this model selection. The models in bold indicate the best model for each response variable. **u**, **Q**, and **R** indicate matrices of the long-term mean population growth rates of four zooplankton groups, and the variance-covariance of the process error and the observation error with a multivariate normal distribution. AICb and ΔAICb indicate bootstrapped Akaike information criterion scores and differences in AICb scores from the best model.

| model | **u** | **Q** | **R** | AICb | ΔAICb | log-likelihood |
| --- | --- | --- | --- | --- | --- | --- |
| m0 | equal | diagonal and equal | diagonal and equal | 2165.6 | 183.0 | -1030.7 |
| m1 | unequal | diagonal and equal | diagonal and equal | 2188.2 | 205.5 | -1029.4 |
| m2 | unequal | diagonal and unequal | diagonal and equal | 2192.4 | 209.8 | -1018.6 |
| m3 | unequal | unconstrained | diagonal and equal | 2029.4 | 46.7 | -851.1 |
| m4 | unequal | diagonal and equal | diagonal and unequal | 2195.5 | 212.9 | -977.8 |
| m5 | unequal | diagonal and unequal | diagonal and unequal | 2167.5 | 184.9 | -949.2 |
| **m6** | **unequal** | **unconstrained** | **diagonal and unequal** | **1982.6** | **0.0** | **-768.3** |
| m7 | unequal | diagonal and equal | grouped by species | 2160.0 | 177.4 | -1011.9 |
| m8 | unequal | diagonal and unequal | grouped by species | 2138.7 | 156.1 | -988.2 |
| m9 | unequal | unconstrained | grouped by species | 2000.7 | 18.0 | -832.4 |
| m10 | unequal | diagonal and equal | grouped by subbasin | 2164.5 | 181.9 | -1015.3 |
| m11 | unequal | diagonal and unequal | grouped by subbasin | 2167.2 | 184.6 | -1003.6 |
| m12 | unequal | unconstrained | grouped by subbasin | 2015.9 | 33.3 | -826.5 |

**Figure S1.**
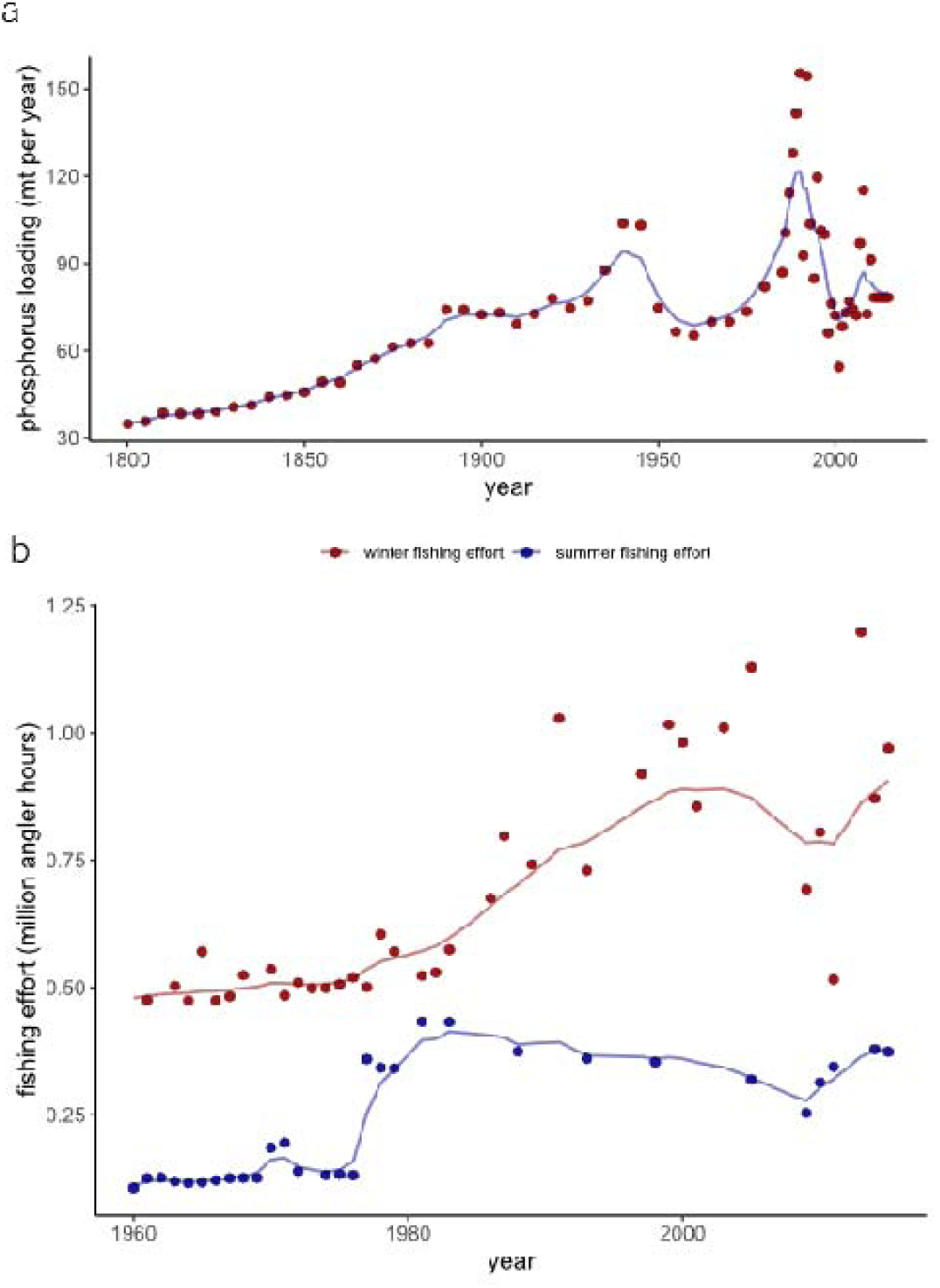
Time series of estimated historical (a) phosphorus loading (1800–2014) and (b) winter and summer fishing effort (1960–2015) in Lake Simcoe. Solid circles indicate data and lines indicate estimates from autoregressive state-space time series models.

## Notes

### Competing Interest Statement

The authors have declared no competing interest.

